# Targeting serine hydroxymethyltransferases 1 and 2 for T-cell acute lymphoblastic leukemia therapy

**DOI:** 10.1101/2020.02.06.936286

**Authors:** Yana Pikman, Nicole Ocasio-Martinez, Gabriela Alexe, Samuel Kitara, Frances F. Diehl, Amanda L. Robichaud, Amy Saur Conway, Angela Su, Jun Qi, Giovanni Roti, Caroline A. Lewis, Alexandre Puissant, Matthew G. Vander Heiden, Kimberly Stegmaier

## Abstract

Despite progress in the treatment of acute lymphoblastic leukemia (ALL), T-cell ALL (T-ALL) has limited treatment options particularly in the setting of relapsed/refractory disease. Using an unbiased genome-scale CRISPR-Cas9 screen we sought to identify pathway dependencies for T-ALL which could be harnessed for therapy development. Disruption of the one-carbon folate, purine and pyrimidine pathways scored as the top metabolic pathways required for T-ALL proliferation. We used a recently developed inhibitor of SHMT1 and SHMT2, RZ-2994, to characterize the effect of inhibiting these enzymes of the one-carbon folate pathway in T-ALL and found that T-ALL cell lines were differentially sensitive to RZ-2994, with a S/G2 cell cycle arrest. The effects of SHMT1/2 inhibition were rescued by formate supplementation. Loss of both SHMT1 and SHMT2 was necessary for impaired growth and cell cycle arrest, with suppression of both SHMT1 and SHMT2 impairing leukemia progression *in vivo*. RZ-2994 decreased leukemia burden *in vivo* and remained effective in the setting of methotrexate resistance *in vitro*. This study highlights the significance of the one-carbon folate pathway in T-ALL and supports further development of SHMT inhibitors for treatment of T-ALL and other cancers.

## Introduction

Metabolic reprogramming is a hallmark of cancer, as cells alter their metabolism to support rapid cell growth and proliferation. As early as the 1920s, Otto Warburg observed that tumor cells consume glucose at a high rate and do fermentation even in the presence of oxygen^1^. Since then drugs targeting metabolism transformed the treatment of certain cancers. In the 1940s, the discovery and application of aminopterin, which was found later to target dihydrofolate reductase (DHFR), a cytoplasmic enzyme involved in one-carbon folate metabolism, yielded the first remission in a child with acute lymphoblastic leukemia (ALL)^2^. Other folate derivatives, such as methotrexate, were later developed. More recently, drugs such as 5-fluorouracil and pemetrexed that target thymidylate synthetase, another enzyme that utilizes folate-derived one-carbon units, were found to be effective therapies for some cancers^3^.

T-cell acute lymphoblastic leukemia (T-ALL) is a highly aggressive cancer characterized by rapid proliferation of early lymphoid cells with immature T-cell surface markers. Early T-cell precursors express the NOTCH1 receptor and rely on high levels of NOTCH1 ligand Δ-like 4 expressed on the surface of thymic epithelial cells during T-cell development^4,5^. High NOTCH1 signaling and expression of the NOTCH1 receptor are thus instructive toward T-cell lineage development. As T-cells progress through thymic development, T cell receptor (TCR) rearrangements are highly coordinated with NOTCH1 signaling and times of rapid cell proliferation in the thymus. Moreover, T-cell activation and development rely on a number of other coordinated pathways including one-carbon folate and specialized mitochondrial proliferation^6^. T-ALL originates as a result of accumulation of mutations that affect cell growth, proliferation and differentiation during this highly coordinated proliferative process. Over 70% of T-ALL, for example, have mutations affecting *NOTCH1* or the NOTCH1 signaling pathway, such as *FBXW7*, highlighting the early normal proliferative signal gone awry^7^.

In addition to NOTCH1, MYC signaling is essential for T-ALL pathogenesis, and MYC is activated by NOTCH in this disease^8^. MYC is a master regulator of cell proliferation, affecting regulation of immortality, cell cycle progression, genetic instability, apoptosis and metabolism^9^. In cancer, pathologic activation of MYC commonly plays a key role in disease pathogenesis. MYC stimulates expression of many mitochondrial genes that are encoded in the nucleus and even regulates mitochondrial biogenesis itself^10^. MYC has been implicated in controlling the one-carbon folate pathway, especially in the presence of hypoxia^11,12^. In the context of acute myeloid leukemia (AML), we have previously shown that MYC binds at promoter sites of enzymes of the one-carbon folate pathway, such as SHMT2, MTHFD2 and MTHFD1L^13^.

While cure rates for pediatric ALL have improved dramatically over the last several decades, leukemia remains the second leading cause of cancer-related death in children^14^. Adult patients with ALL continue to have poor prognosis even with intensive therapy. T-ALL, comprising 15-20% of ALL, is associated with early relapses and is more likely to be refractory to treatment in the relapse setting^15,16^. Moreover, while there is excitement surrounding immune-mediated therapies for B-cell acute lymphoblastic leukemia (B-ALL), including antibody-based approaches and CAR T-cell therapies, these approaches are not available for patients with T-ALL. Thus, more effective therapies are needed for patients with T-ALL, particularly those with relapsed or refractory disease.

Given the role of the one-carbon folate pathway in cancer there is an interest in developing novel inhibitors of this pathway. Inhibition of the one-carbon folate pathway has already yielded a number of highly active drugs, such as methotrexate (targeting DHFR), 5-fluorouracil and pemetrexed (targeting thymidylate synthetase and DHFR), gemcitabine (a deoxycytidine analog), and mercaptopurine (a PRPP aminotransferase inhibitor inhibiting purine nucleotide synthesis). Over the last several years, there has been an increase in the development of inhibitors of the de novo purine and pyrimidine synthesis pathways, with drugs such as DHODH inhibitors showing promise in early phase clinical trials^17^. Inhibitors of plasmodial serine hydroxymethyltransferase (SHMT) have been developed for malaria treatment. These pyrazolopyran-based ligands have been optimized for *in vitro* activity against SHMT enzymes, though with limited *in vivo* activity^18^. Further optimization for SHMT activity has led to an *in vitro* selective SHMT1/2 inhibitor, though this has not been tested *in vivo*^19^.

Given the clinical need for the treatment of T-ALL, we used an unbiased CRISPR-Cas9 genome-scale screen to identify specific pathway dependencies for T-ALL. The one-carbon folate, purine and pyrimidine pathways scored as the top metabolic pathways in this analysis. Using a combination of small molecule inhibitors and genetic suppression of SHMT, we validate the combined repression of SHMT1 and SHMT2 as a candidate therapeutic approach for T-ALL.

## Methods

### Cell Culture, Cell Viability and Flow Cytometry Assays

PF382, RPMI8402, KOPTK1 and HSB2 cell lines were obtained from Dr. Jon Aster, and identity verified using STR profiling. All cell lines were maintained in RPMI 1640 (Cellgro) supplemented with 1% penicillin/streptomycin (PS)(Cellgro) and 10% FBS (Sigma-Aldrich) at 37°C with 5% CO_2_. Viability was evaluated using the CellTiter-Glo Luminescent Cell Viability Assay (Promega) after the indicated days of exposure to the specific drug or combination of drugs. Luminescence was measured using FLUOstar Omega from BMG Labtech. The IC_50_ values were determined using Prism GraphPad version 8 software.

For cell cycle analysis, T-ALL cells were harvested at the indicated time points, washed and fixed in ethanol and then re-suspended in 49 μg/mL propidium iodide (Sigma-Aldrich) and 100 μg/mL of RNase A (Qiagen). Cell death was assessed using flow cytometric analysis of Annexin V and propidium iodide staining according to the manufacturer’s instructions (eBioscience). Samples were analyzed on a FACSCanto analyzer (BD Biosciences). Data analysis was completed using Flowjo software.

### Compounds

RZ-2994 was initially obtained from Raze Therapeutics. After the structure was published^6^ and Raze Therapeutics was no longer producing it, RZ-2994 was synthesized by Medicilon. Identity was confirmed independently by LC-MS and NMR performed by Dr. Jun Qi (DFCI). Methotrexate was purchased from Sigma Aldrich.

### CRISPR-Cas9 Screening

The CRISPR-Cas9 screen was performed on the Avana library containing 73,372 guides for 18,333 genes, with an average of 4 guides per gene. The analysis was done on the 19Q4 version of the gene effect Avana data processed with the CERES algorithm^20^, publicly available on the Depmap portal https://depmap.org/portal/. This dataset contains 689 cell lines, including 3 T-cell ALL lines: KOPTK1, PF382 and HSB2, and 73 other hematopoietic cell lines.

Initially, cancer cell lines were transduced with Cas9 using a lentiviral system. Cell lines that met quality criteria, including acceptable Cas9 measured ability to knockout transduced GFP, appropriate growth properties and other parameters, were then screened with the Avana library. A pool of guides was transduced into a population of cells. The cells were cultured for 21 days *in vitro*, and at the end of the assay, barcodes for each guide were sequenced for each cell line in replicate.

The sgRNA read count data were deconvoluted from sequence reads by using the PoolQ public software (https://portals.broadinstitute.org/gpp/public/software/poolq). A series of quality control pre-processing steps was performed to remove samples with poor replicate reproducibility, as well as guides that have low representation in the initial plasmid pool, as described by Dempster et al^21^. The raw read counts were summed up by replicate and guide and the log2-fold-change from pDNA counts for each replicate was computed. The sgRNAs with suspected off-target activity and the guides with pDNA counts less than one millionth of the pDNA pool were removed. The replicates that failed fingerprinting and the replicates with less than 15 million reads were removed and then the replicate read counts were scaled to 1 million total reads per replicate. The replicates with the null-normalized mean difference (NNMD) greater then −1.0 were filtered out. Also removed were those replicates that did not have a sufficiently high Pearson coefficient (> 0.61) with at least one other replicate for the line when looking at genes with the highest variance (top 3%) in gene effect across cell lines. Then NNMD was computed again for each cell line after averaging remaining replicates, and the cell lines with NNMD > −1.0 were filtered out.

For quality control and normalization, exogenously defined nonessential genes^22^ were used as negative controls, and common essential genes^23,24^ were used as positive controls. The gene level dependency scores were inferred by running the computational tool CERES^20^. CERES was developed to computationally correct the copy-number effect and to infer true underlying effect of gene knockout. CERES models the observed normalized log-fold change for each sgRNA and cell line as the linear combination of gene-knockout and copy-number effects with coefficients giving the guide activities. Copy-number effects are fit with a linear piece-wise model in each cell line. Once all parameters have been fit, the inferred gene scores and guide activity scores are extracted and reported.

The CERES gene dependency data was further scaled to the −1 value of the median of common essential genes in each cell line, and then transformed into z-scores so that each gene had mean = 0 and variance = 1. Next the first five principal components of the resulting data were removed, the prior means of genes were restored, and the data were scaled again so the median of common essentials in each cell lines was -1. The pan-dependent genes were identified as those genes for whom 90% of cell lines rank the gene above a dependency cutoff determined from the central minimum in a histogram of gene ranks in their 90th percentile least dependent line. For each CERES gene score, the probability that the score represents a true dependency or not was inferred based on the expectation-maximization algorithm.

The differential dependency gene level scores for the T-ALL lineage were determined for T-ALL vs. all other non T-ALL cell lines, and also for T-ALL vs. all other non T-ALL hematopoietic cell lines, in order to eliminate the bias induced by the hematopoietic lineage itself. The analysis was performed based on the empirical Bayes (eBayes) statistics available from the limma package^25^ (Bioconductor v3.10 https://www.bioconductor.org/packages/release/bioc/html/limma.html) with the significance cutoffs: abs(size effect) ≥ 0.3, P-value ≤ 0.05, adjusted P-value ≤ 0.10.

### Vectors and Constructs

shRNA constructs targeting SHMT1 and SHMT2 were designed and delivered via a LT3 GEPIR (SHMT1) or REVIR(SHMT2) vector as previously described^26^. Hairpin sequences are listed in Supplementary Table 1. For virus production, 12 μg of the above vector with 6 μg pCMV8.9 and pCMV-VSVG packaging vectors were transfected into the 293 packaging cell line using X-tremeGENE 9 (Roche), and the resulting viral supernatants were harvested as previously described^13^. sgRNA constructs were designed using the Broad Institute’s shRNA designer tool; sequences are listed in Supplementary Table 1.

### RNASeq

KOPTK1 cells were grown in the presence of 2 µM RZ-2994 vs DMSO and cells collected at 24 and 72 hours of treatment. Three samples were collected per treatment condition per time point. RNA was extracted from cells with an RNeasy Kit (Qiagen) and was sequenced using Illumina TruSeq strand specific library. Quality control tests for the 75 bp single-end mapped reads were performed using the FASTQC software (www.bioinformatics.babraham.ac.uk/projects/fastqc/). The reads were aligned to the GRCh37/hg19 human genes by using STAR v2.7-2b^27^. Quality control tests for the aligned reads and for replicate consistency were performed by using the qualimap v2.2.1^28^ and the SARTools^29^ pipelines. The RNA-Seq data for this study is available for download from the Gene Expression Omnibus (GEO) repository https://www.ncbi.nlm.nih.gov/geo/ (GSE143176) upon manuscript publication.

Gene level reads and gene level expression estimated as log2(1+TPM) scores – where TPM stands for Transcripts Per Million - were computed using the Feature Counts method implemented in the Bioconductor v3.10 RSubread package^30^. The overall significance of the differential expression between the control (DMSO) and treatment (RZ-2994) phenotypes at Day 1 and separately at Day 3, was estimated by using the apeglm method^31^ available from the DESeq2 library^32^ (Bioconductor v3.10) with the standard significance cut-offs abs(shrinkage fold change) ≥ 1.5, adjusted P-value ≤ 0.10.

### Gene Set Enrichment Analysis (GSEA) for T-ALL Dependencies

The GSEA v4.0.3 software^33,34^ was utilized to identify the Kyoto Encyclopedia of Genes and Genomes (KEGG) canonical pathways that have a significant overlap with the genes showing a differential dependency for the T-ALL vs. non T-ALL cell lines and separately, for the T-ALL vs. non T-ALL hematopoietic cell lines in the Avana 19Q4 dependency data. First, the hg19 genes were ranked in decreasing order based on the T-ALL differential dependency scores. The goal of GSEA was to identify the pathways that are distributed at the top or at the bottom of the ranked list of genes. For this purpose, the Pre-Rank GSEA module was run across the collection of 186 KEGG pathways available in the MSigDB v7.0 database^33,35,36^ with the significance cut-offs nominal P-value ≤ 0.10 and FDR ≤ 0.25 for the Kolmogorov-Smirnov enrichment test. The significantly enriched pathways with the Normalized Enrichment Score (NES) ≤ −1.5 were annotated for “Depletion” in T-ALL and those with NES ≥ 1.5 were annotated for “Proliferation” in T-ALL. The KEGG pathways identified as significantly enriched in dependency genes for T-ALL were further manually annotated as related to the amino-acid metabolic functional category.

### Single-sample Gene Set Enrichment Analysis for T-ALL Dependencies

A single-sample GSEA (ssGSEA) analysis^37,38^ was performed on the CERES dependency data across the collection of 186 KEGG pathways available from the MSigDB v7.0 database to further analyze the functional association of the amino-acid metabolic pathways with T-ALL dependencies.

ssGSEA is a variant of the GSEA method that assigns to each individual sample, represented as a ranked list of genes, an Enrichment Score (ES) with respect to each gene set in a given collection of pathways. The ssGSEA ES is calculated as a running sum statistic by walking down across the ranked list of genes, increasing the sum when encountering genes in the gene set and decreasing it when encountering genes not in the gene set. The significance of the ES is estimated based on a permutation P-value and adjusted for multiple hypotheses testing through FDR. A positive ES denotes a significant overlap of the signature gene set with groups of genes at the top of the ranked list, while a negative ES denotes a significant overlap of the signature gene set with groups of genes at the bottom of the ranked list.

For each sample, the ES is further transformed into a Z-score by subtracting the average of the ES’s assigned to all other samples and by dividing the result to their standard deviation. While GSEA generates a gene set’s enrichment score with respect to phenotypic differences across a collection of samples within a dataset, ssGSEA calculates a separate enrichment score for each pairing of sample and gene set, independent of phenotype labeling. In this manner, ssGSEA transforms a single sample’s dependency profile to a gene set enrichment profile. A gene set’s enrichment score represents the activity level of the biological process in which the gene set’s members are coordinately scoring up or down. The ssGSEA gene set representation has an unsupervised biological interpretability and can be further analyzed with statistical and machine learning methods.

### Metabolite Profiling and Analysis

For metabolite extraction, 1 million cells per condition were pelleted and washed with ice cold saline. Cell pellets were resuspended in 1 mL of 80% methanol solution containing 500 nM internal standards (Metabolomics Amino Acid Mix, Cambridge Isotope Laboratories Inc.). Samples were vortexed at 4 degrees, followed by centrifugation at 4C for 10 minutes. Supernatant was transferred to a new tube, and samples dried using a Speedvac. Samples were collected in triplicate.

Dried cell extracts were resuspended in 50 μL HPLC grade water. LC-MS analysis was performed using a QExactive orbitrap mass spectrometer using an Ion Max source and heated electro-spray ionization (HESI) probe coupled to a Dionex Ultimate 3000 UPLC system (Thermo Fisher Scientific). External mass calibration was performed every 7 days. Typically, samples were separated by chromatography by injecting 2 μL of sample on a SeQuant ZIC-pHILIC 2.1 mm × 150 mm (5 μm particle size) column. Samples were run at multiple dilutions to ensure linearity of all metabolites measured. Flow rate was set to 150 mL/min. and temperatures were set to 25C for the column compartment and 4C for the autosampler tray. Mobile phase A was 20 mM ammonium carbonate, 0.1% ammonium hydroxide. Mobile phase B was 100% acetonitrile. The chromatographic gradient was: 0–20 min.: linear gradient from 80% to 20% mobile phase B; 20–20.5 min.: linear gradient from 20% to 80% mobile phase B; 20.5 to 28 min.: hold at 80% mobile phase B. The mass spectrometer was operated in full scan, polarity-switching mode and the spray voltage was set to 3.0 kV, the heated capillary held at 275C, and the HESI probe was held at 350C. The sheath gas flow rate was 40 units, the auxiliary gas flow was 15 units and the sweep gas flow was one unit. The MS data acquisition was performed in a range of 70–1000 m/z, and an additional narrow-range scan (220-700 m/z) was included in negative mode to enhance the detection of nucleotides. The resolution was set at 70,000, the AGC target at 1×106, and the maximum injection time at 20 msec. Relative quantitation of polar metabolites was performed with TraceFinder 4.1™ (Thermo Fisher Scientific) using a 5 ppm mass tolerance and referencing an in-house library of chemical standards. Peak areas were normalized to internal standards and cell number.

### Immunoblotting

Cells were lysed in Cell Signaling Lysis Buffer (Cell Signaling Technology) as previously reported^13^ and resolved by gel electrophoresis using Novex 4-12% Bis-Tris Gels (Invitrogen), transferred to a nitrocellulose membrane (Bio-Rad) and blocked for one hour in 5% BSA (Sigma). Blots were incubated in primary antibody to SHMT1 (Cell Signaling, #80715), SHMT2 (Cell Signaling, #12762) or Vinculin (Cell Signaling, #13901), followed by the secondary antibodies anti-rabbit HRP (Amersham) or anti-mouse HRP (Amersham). Bound antibody was detected using the Western Lightning Chemiluminescence Reagent (Perkin Elmer).

### *In Vivo* Studies

For genetic inhibition studies, RMPI8402 cells were infected with lentivirus targeting renilla (CTL), SHMT1, SHMT2 or the combination, and cells selected. 750,000 cells were injected via the tail vein into 8-week-old, female NSG mice (The Jackson Laboratory). Disease burden was followed using peripheral blood hCD45. At the time of disease detection of at least 1% human cells, mice were switched to receive doxycycline 2000 ppm chow. Mice were treated for 9 days prior to disease assessment.

For the RZ-2994 therapeutic study, 500,000 cells RPMI8402 luciferized cells were injected via the tail vein into 8-week-old, female NSG mice (The Jackson Laboratory). Leukemia burden was assessed using non-invasive bioluminescence imaging by injecting mice intraperitoneally with 75 mg/kg d-Luciferin (Promega), anesthetizing them with 2–3% isoflurane, and imaging them on an IVIS Spectrum (Caliper Life Sciences). A standardized region of interest (ROI) encompassing the entire mouse was used to determine total body bioluminescence, with data expressed as photons/s/ROI (ph/s/ROI). Once detectable bioluminescence was achieved, the mice were separated into two treatment cohorts (RZ-2994 and vehicle), 7 mice per cohort and treatment initiated. Mice were treated with RZ-2994 at 100 mg/kg IP daily for 14 days. There was no blinding of the person treating the mice to the treatment. Sample size was calculated to have 80% power to detect 1.75 SD difference between the two groups using a two-sided t-test with α = 0.05. All animal studies were conducted under the auspices of protocols approved by the Dana-Farber Cancer Institute Animal Care and Use Committee.

### Drug Interaction Analysis

The expected dose-inhibitory fraction relationships for the combination therapy of RZ-2994 and methotrexate were assessed using the Bliss independence model^39,40^. The Bliss Independence model is based on the principle that drug effects are outcomes of probabilistic processes and compares the effect resulting from the combination of two drugs directly to the effects of its individual components. The model computes a quantitative measure called excess over Bliss (*eob*). Positive *eob* values are indicative of synergistic interaction whereas negative *eob* values are indicative of antagonistic behavior. Null *eob* values indicate additive effect.

### Statistical Analysis

Statistical significance was determined by two-tailed t test or Mann-Whitney test for pair-wise comparison of groups, as indicated. Statistical calculations were performed using Prism GraphPad version 8 software.

## Results

### One-carbon folate metabolism is a dependency in T-ALL

To identify selective pathway dependencies for T-ALL we used CRISPR-Cas9 whole genome screening data of 689 cancer cell lines from the Broad Institute Dependency Map project^41^. Three T-ALL cell lines were included in the screen, in addition to 73 other hematopoietic cell lines and 613 cell lines derived from solid tumors. Gene set enrichment analysis (GSEA) was performed against 186 KEGG pathways to identify top negatively enriched pathways for T-ALL compared to other cancers. Inhibition of these pathways would be predicted to be more therapeutically effective in T-ALL compared to other tumors. The one-carbon folate, purine and pyrimidine KEGG pathways scored among the top dependencies in T-ALL versus all other cancer cell lines (Fig. 1A) and when compared to other hematopoietic cell lines (Fig. 1B). As a positive control, known T-ALL pathway dependencies, such as NOTCH signaling and T-cell receptor signaling, were also among the top 10 KEGG scoring pathways (Supplementary Tables 2A and 2B). Focusing on these pathways separately, each was a significant dependency in T-ALL (Fig.1C and Supplementary Fig. 1). To further validate this finding in a primary T-ALL dataset, we performed single sample GSEA (ssGSEA) on a large human primary ALL gene expression data set (St. Jude, 575 ALL samples, including 84 T-ALL samples^42^) for enrichment across the collection of 186 canonical KEGG pathways. T-ALL samples showed significantly increased expression of genes involved in the one-carbon folate, purine and pyrimidine metabolism pathways, compared to B-ALL samples in this data set (Fig. 1D). We validated this finding in a second gene expression data set with 107 primary ALL samples, including 15 T-ALL samples (GSE13351, Supplementary Fig. 2A)^43^. Both data sets showed a significant enrichment of the one-carbon folate pathway associated with the T-ALL vs. other non-T-ALL samples (P ≤ 0.0001, Supplementary Fig. 2B and 2C).

**Figure 1:**
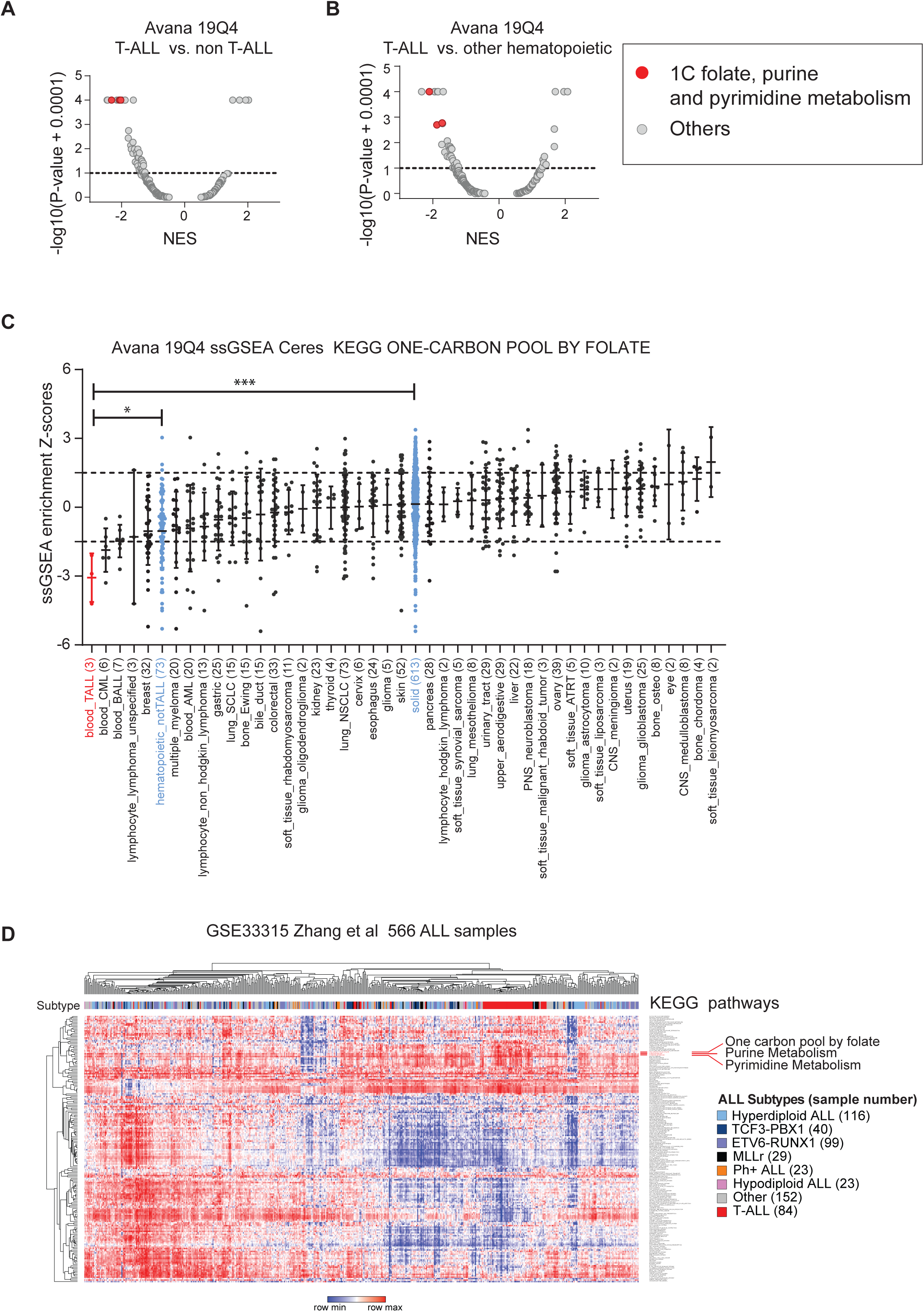
One-carbon folate metabolism is a dependency in T-ALL. Volcano plots showing enrichment of KEGG pathways for 689 cell lines in the Avana 19Q4 dataset. The KEGG one-carbon pool by folate, purine metabolism and pyrimidine metabolism pathways scored as most depleted in the T-ALL lineage (n=3) compared to all other cell lines (n=686) (A) (P = 0.0003, Mann-Whitney test) and compared to other hematopoietic cell lines (n=73) (B) (P = 0.025, Mann-Whitney test). Normalized enrichment score (NES) shown on X-axis. C) Graph showing the distribution of the ssGSEA Z-scores for the one-carbon pool by folate pathway across cancer cell lineages represented in the Avana 19Q4 data set. The one-carbon pool by folate pathway is significantly enriched in T-ALL vs non-T-ALL hematopoietic (*P < 0.05, Mann-Whitney test) and T-ALL vs solid tumor (***P < 0.001, Mann-Whitney test) cell lines. D) Heatmap of ssGSEA projection for the primary ALL sample data set GSE33315 from St. Jude on the collection of KEGG canonical pathways. T-ALL samples are highlighted in red.

Inhibitors of the one-carbon folate pathway, such as methotrexate and mercaptopurine, have formed the backbone of ALL therapy. Given the selective dependency on the one-carbon folate pathway in T-ALL, we tested RZ-2994^6^ (also known as SHIN1^19^) as a novel inhibitor of this pathway. RZ-2994 is an inhibitor of the cytoplasmic SHMT1 and mitochondrial SHMT2 serine hydoxymethyltransferases^19,44^. For comparison to other hematopoietic cell lines, we tested AML, B-ALL and T-ALL cells for sensitivity to RZ-2994. T-ALL was sensitive to RZ-2994 compared to other acute leukemia cell lines, with an average IC_50_ of 1.6 μM (Fig. 2A). Treatment of 4 T-ALL cell lines resulted in accumulation of cells in the S and G2 cell cycle phases with minimal apoptosis (Fig. 2B, Supplementary Fig. 3 and Fig. 2C). We next evaluated the effects of SHMT inhibition on gene expression. We treated the KOPTK1 cell line with RZ-2994 for 1 and 3 days and performed RNA sequencing analysis. RZ-2994 treatment resulted in changes in pathways associated with amino acid metabolism (Fig. 2D and Supplementary Fig. 4A), MYC targets (Fig. 2E and Supplementary Fig. 4B) and associated with cell cycle arrest (Fig. 2F and Supplementary Fig. 4C). These changes were more pronounced after 3 days of treatment.

**Figure 2:**
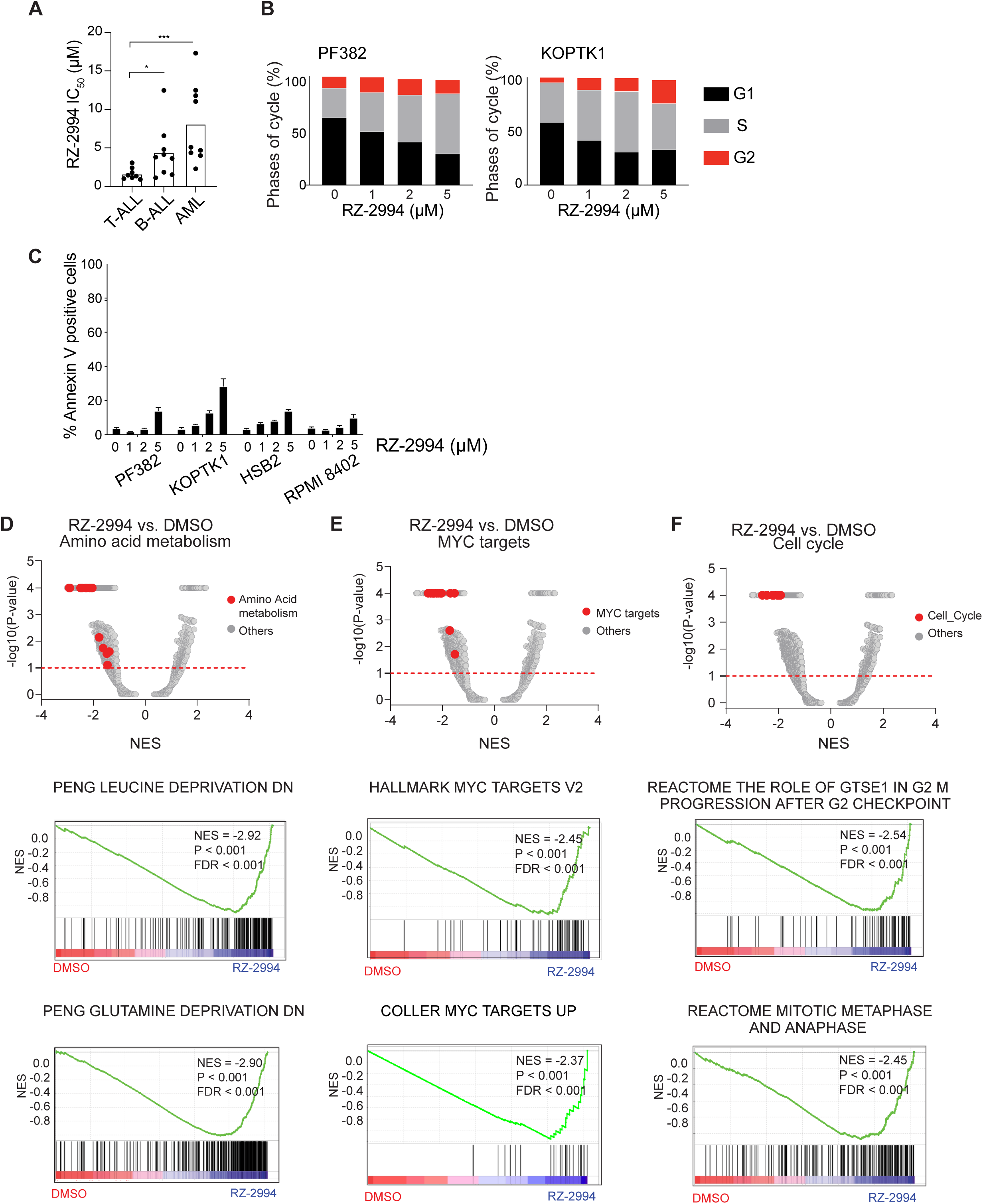
Enzymatic inhibition of SHMT1 and SHMT2 results in T-ALL arrest and gene expression changes. A) T-ALL (n=8), B-ALL (n=9) and AML (n=9) cell lines were treated with RZ-2994 in a range of concentrations, in quadruplicate for 6 days. Bar graph showing the average IC_50_ per lineage, with each dot representing the IC_50_ in a cell line. *P<0.05 and ***P<0.001 using a Mann-Whitney test. B) Cell cycle analysis in T-ALL cells treated with increasing concentrations of RZ-2994. C) Bar graph showing percent Annexin V positive cells with increasing concentrations of RZ-2994 in T-ALL cell lines. Shown are the mean ± standard deviation (SD) of 3 replicates. RNAseq was performed for the KOPTK1 cell line treated with RZ-2994. Volcano plots showing quantitative comparison of gene sets from MSigDB v7.0 using ssGSEA. Volcano plots compare DMSO versus RZ-2994 after 3 days of treatment. All datasets above the dashed red line have P-value ≤ 0.05. Gene expression changes associated with 3-day RZ-2994 treatment show enrichment for D) amino acid metabolism, E) MYC targets and F) cell cycle pathways. Top scoring GSEA plots are shown below the associated volcano plots.

Inhibition of SHMT1 and SHMT2 impairs glycine and formate synthesis, which in turn can impede nucleotide production^19,44^. We performed metabolite profiling of PF382, KOPTK1 and RPMI8402 cell lines treated with RZ-2994 for 3 days and observed changes in intermediates that involve the one-carbon folate pathway (Supplementary Fig. 5). We focused on metabolic changes that were common among the three cell lines as those are more likely to contribute to the RZ-2994-related effect on cell growth. Consistent with SHMT1 and SHMT2 inhibition, we found glycine levels were decreased with an increase in serine levels. We also observed increases in the purine precursors AICAR and GAR (Fig. 3A), which are upstream of steps of the purine synthesis where one-carbon units are incorporated. There was also a decrease in ATP and dTTP. Thymidylate synthase requires 5,10-methylenetetrahydrofolate to synthesize dTMP from dUMP. Consistent with possible depletion of THF by SHMT1 and SHMT2 inhibition, there was an increase in dUMP (Fig. 3A). Addition of 1 mM formate rescued the proliferation of T-ALL cell lines in the presence of RZ-2994 (Fig. 3B), including rescue of the cell cycle arrest (Fig. 3C). Cells cultured in the presence of formate supplementation were no longer sensitive to the effects of RZ-2994 (Fig. 3D).

**Figure 3:**
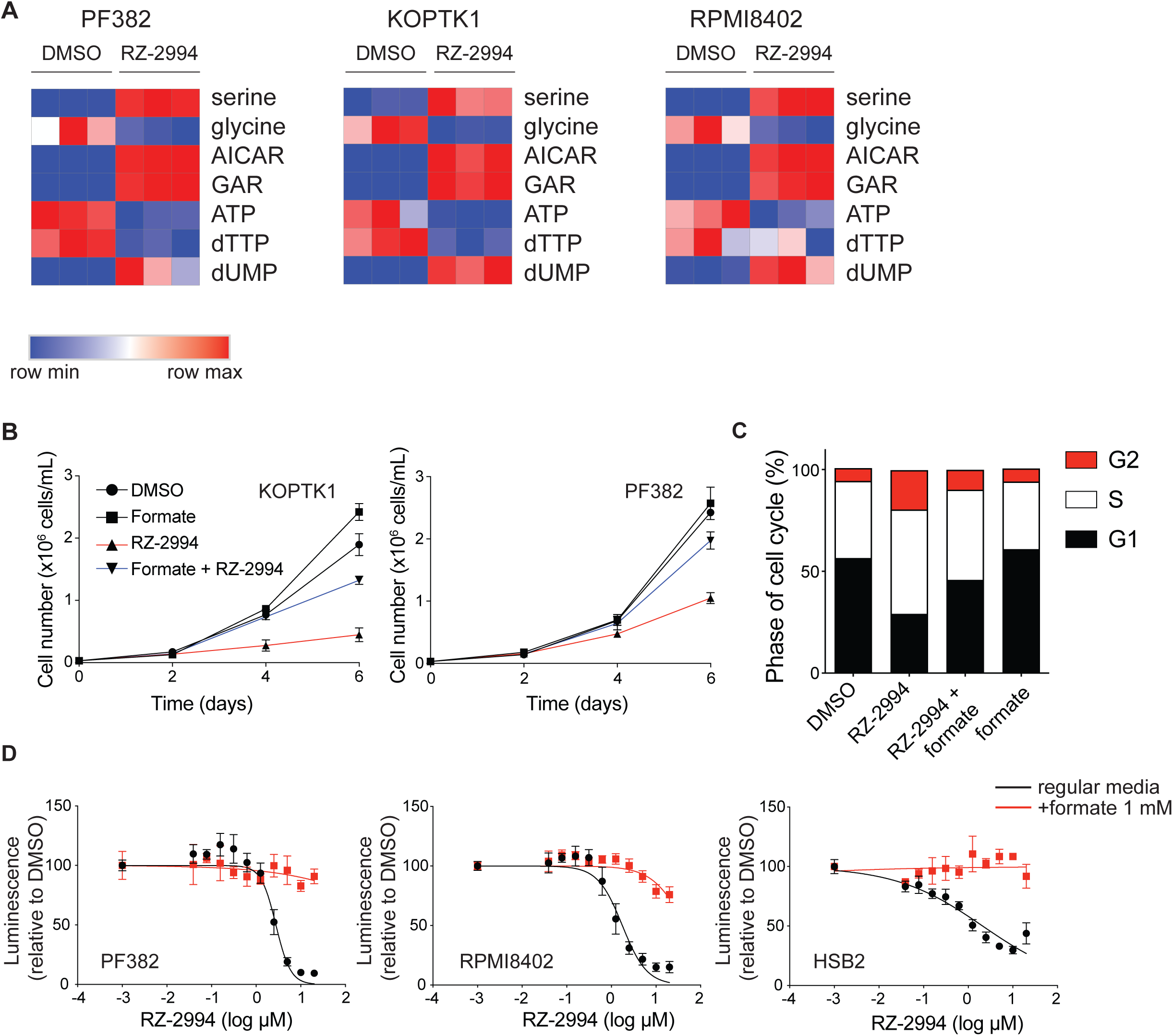
RZ-2994 causes metabolic changes in T-ALL, and its antiproliferative effects can be rescued with formate supplementation. A) Heatmaps showing changes in metabolites associated with one-carbon folate metabolism following treatment with RZ-2994 in 3 cell lines. Cell lines were treated with 2 µM RZ-2994 for 3 days, metabolites extracted and profiled using LC-MS. Raw peak areas were normalized to internal standards. Heatmap shows normalization of the relative metabolite abundance per metabolite. B) RZ-2994 leads to a decreased cell growth in T-ALL cell lines, and this growth defect can be rescued with supplementation of 1 mM formate. Graphs depict cell number as measured by trypan blue exclusion. Shown are the means ±SD of 3 replicates. C) Cell cycle analysis in T-ALL cells treated with DMSO, RZ-2994 (2 µM), formate (1 mM) or the combination of RZ-2994 with formate. D) T-ALL cell lines were grown in a range of RZ-2994 concentrations, in regular media or supplemented with 1 mM formate, and viability evaluated at day 6 by an ATP-based assay as the percentage of viable cells relative to a DMSO control. Shown are the mean ± standard deviation (SD) of 4 replicates.

### Loss of both SHMT1 and SHMT2 is necessary to impair proliferation of T-ALL

Enzymes of the mitochondrial one-carbon folate pathway are highly expressed in cancer,^45,46^ and their expression has been associated with poor survival^46^. We have previously shown MTHFD2 and enzymes of the one-carbon folate pathway to be highly expressed in AML, with inhibition of MTHFD2 leading to a decrease in AML viability *in vitro* and *in vivo*^13^. Redundancy of SHMT1 and SHMT2 enzymes has been shown in HEK293T cells and HCT-116 colon cancer cells, though it is unclear if this occurs in leukemia cells^47^. Given that RZ-2994 inhibits both the cytoplasmic and mitochondrial SHMT enzymes, we evaluated if both need to be inhibited to affect proliferation of T-ALL cells. We used shRNA to knockdown *SHMT1, SHMT2* individually or both genes together. Repression of either *SHMT1* (Fig. 4A) or *SHMT2* (Fig. 4B) was not sufficient to impair cell proliferation. SHMT1 or SHMT2 individually were not dependencies in the CRISPR-Cas9 screening data (Supplementary Fig. 6). Instead, loss of both *SHMT1* and *SHMT2* was necessary for the full anti-proliferative effect and cell cycle arrest (Fig. 4C and 4D). We also used CRISPR-Cas9 to knock out *SHMT1, SHMT2* or the combination, with an anti-proliferative effect observed only upon loss of both *SHMT1* and *SHMT2*, and this effect rescued by formate supplementation (Fig. 4E).

**Figure 4:**
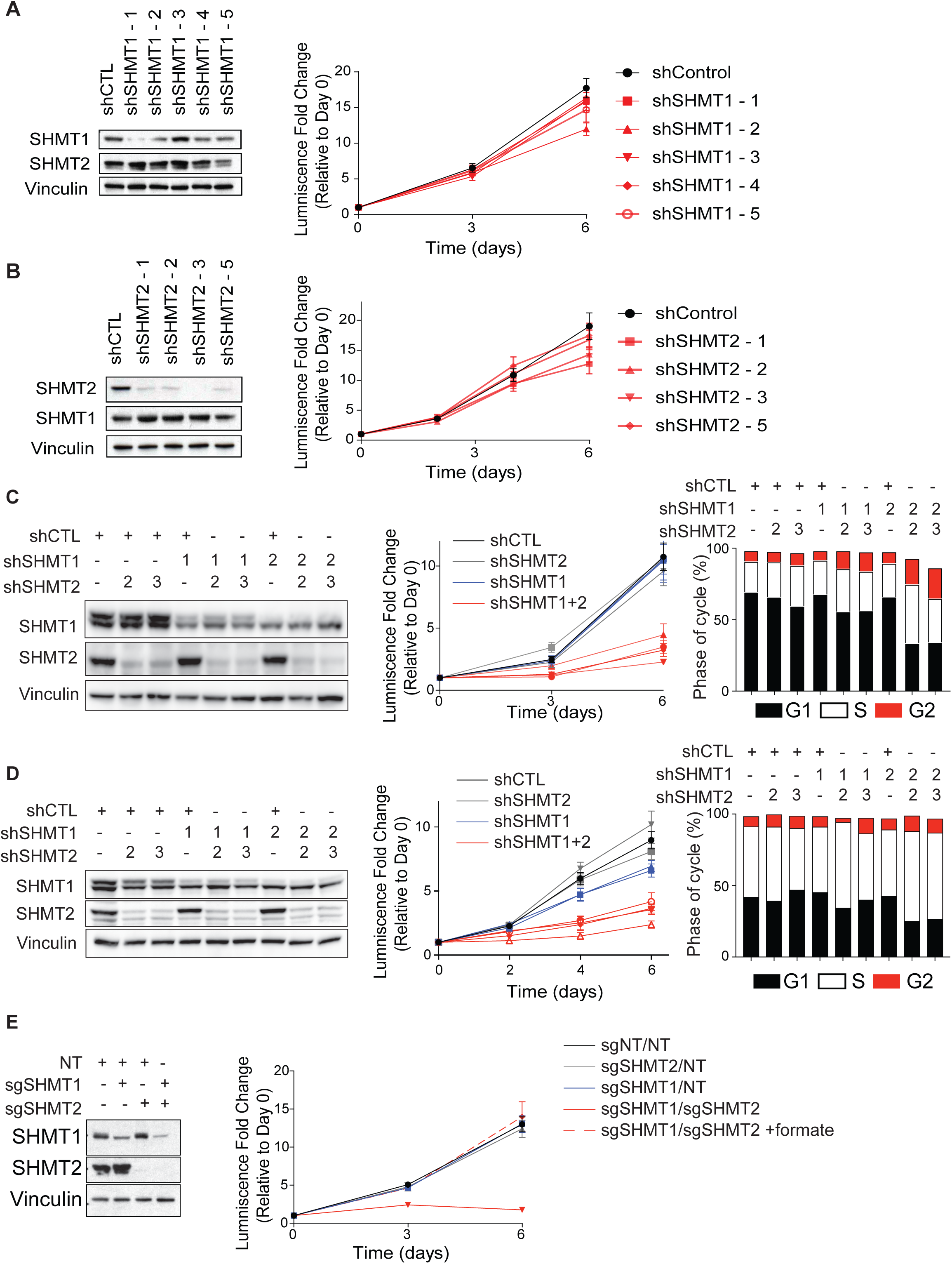
Loss of both SHMT1 and SHMT2 is required for T-ALL cell cycle arrest. A) Western blot evaluating knockdown of SHMT1 in a PF382 cell line with five unique doxycycline-inducible shRNAs (shSHMT1-1, shSHMT1-2, shSHMT1-3, shSHMT1-4 and shSHMT1-5) compared to a control shRNA (shControl). Vinculin is used as a loading control. Cells were grown over the course of 6 days and viability assessed by an ATP-based assay. Graphs depict luminescence fold change per cell line condition relative to Day 0. Shown are the means ±SD of 4 replicates. B) Western blot evaluating knockdown of SHMT2 in the PF382 cell line with four unique doxycycline-inducible shRNAs (shSHMT2-1, shSHMT2-2, shSHMT2-3 and shSHMT2-5) compared to a control shRNA (shControl). Vinculin is used as a loading control. Cells were grown over the course of 6 days and viability assessed by an ATP-based assay. Graphs depict luminescence fold change per cell line condition relative to Day 0. Shown are the means ±SD of 4 replicates. We used a combination of SHMT1 and SHMT2 targeting hairpins to knockdown SHMT1, SHMT2 or both in PF382 cells (C) or RPMI8402 cells (D). Western blot showing knockdown using shSHMT1-1 or shSHMT1-2 (labeled 1 or 2 in the SHMT1 row), or shSHMT2-2 or shSHMT2-3 (labeled 2 or 3 in the SHMT2 row). Addition of shControl vectors shown with +. Cells were grown over the course of 6 days and viability assessed by an ATP-based assay. Graphs depict luminescence fold change per cell line condition relative to Day 0. shSHMT2-2 and shSHMT2-3 are both labeled “shSHMT2” and colored grey. shSHMT1-1 and shSHMT1-2 are both labeled “shSHMT1” and colored blue. Cells with knockdown of both SHMT1 and SHMT2 are shown in red. Shown are the means ±SD of 4 replicates. Bar graph showing cell cycle analysis in cells after inducible shRNA knockdown. E) Western blot showing knockout of *SHMT1, SHMT2* or both using CRISPR guides. Cells were grown over the course of 6 days and viability assessed by an ATP-based assay. Graphs depict luminescence fold change per cell line condition relative to Day 0.

### SHMT inhibition has *in vivo* efficacy in T-ALL

In order to study the effects of SHMT1 and SHMT2 suppression after the development of T-ALL *in vivo*, we deployed a doxycycline-inducible shRNA system directed against SHMT1 and SHMT2 using the two constructs that yielded efficient suppression of these genes *in vitro* (Fig.4D). RPMI8402 cells bearing the doxycycline-inducible shRNA directed against SHMT1, SHMT2 or the combination were injected into NSG mice. Leukemia establishment was confirmed by hCD45 detection in peripheral blood, and then mice were treated with doxycycline for 9 days until disease progression. Cells induced for loss of SHMT1 (or associated control) become GFP+, while induction of SHMT2 (or its control) results in DsRed expression (Fig. 5A). At time of disease evaluation, we selected hCD45+ cells that were GFP+ and dsRed+ by flow cytometry (Fig. 5B). Suppression of both SHMT1 and SHMT2 led to a competitive disadvantage, with a decrease in these double knockdown cells compared to controls (Fig. 5C).

**Figure 5:**
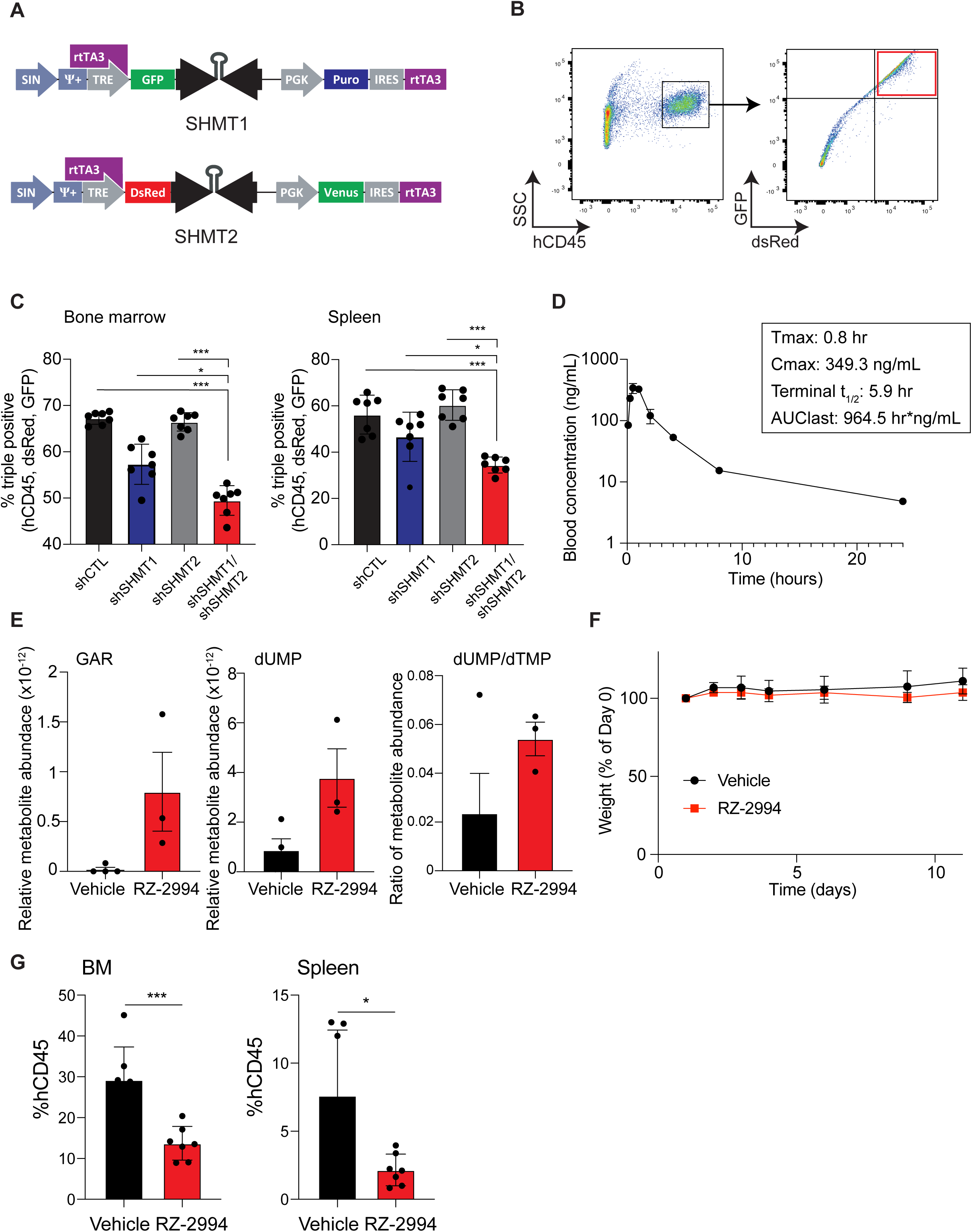
Knockdown and enzymatic inhibition of SHMT1 and SHMT2 are effective for T-ALL therapy *in vivo*. A) Schematic showing SHMT1 and SHMT2 targeting doxycycline inducible constructs which were used for the cell line and mouse experiments. B) Sample flow plot showing schema for cell selection with both SHMT1 and SHMT2 hairpins (or associated controls) for the *in vivo* study. C) Bar graph depicting percent of triple positive (hCD45+, GFP+ and dsRed+) cells in bone marrow and spleen. Shown is average with SD, n=7 per group. *P<0.05, ***P<0.001 using Mann-Whitney test. D) Graph showing blood RZ-2994 concentrations over time after a single dose of 20 mg/kg IP. Shown is the average ± SD, n=3 per group. E) Polar metabolites were extracted from spleens of mice treated with RZ-2994 for 1 week and targeted profiling done using LC-MS. Bar graph shows relative metabolites compared to internal controls. Shown is average with SD, n=4 for vehicle samples, and n=3 for RZ-2994 samples. F) Irradiated NSG mice were injected with RPMI8402-lucNeo cells. After disease was established, mice were treated with RZ-2994. Graph showing weights relative to Day 0 of treatment. Shown is average with SD, n=7 per group. After 2 weeks of treatment, leukemia burden was assessed. G) Bar graph showing percent of hCD45+ cells in bone marrow and spleen after treatment with RZ-2994 100 mg/kg for 2 weeks. *P<0.05, ***P<0.001 using Mann-Whitney test.

RZ-2994 was reported to have limited stability in liver microsome assays^19^, though related pyrazolopyrans have shown modest efficacy *in vivo* for malaria treatment with oral dosing^18^. We thus performed pharmacokinetic analysis of RZ-2994 to assess its bioavailability for use as a tool compound *in vivo*. After injection of RZ-2994 20 mg/kg IP (intraperitoneally), serial levels were measured, with a t_1/2_=5.9 hours and drug levels shown in Figure 5D. We next performed a dose escalation study, where NSG mice were treated with up to 100 mg/kg without toxicity for 1 week. We next conducted a pilot experiment to test whether RZ-2994 treatment results in predicted metabolic changes in an orthotopic RPMI8402 mouse model, with RZ-2994 dosed at 100 mg/kg IP daily. Three mice per group were treated for 1 week and selected metabolites profiled. In line with the *in vitro* data, RZ-2994 led to a trend toward increased GAR and dUMP, and an increase in the dUMP/dTMP ratio, consistent with disrupting one-carbon folate metabolism *in vivo* (Fig. 5E).

We next investigated the *in vivo* efficacy of RZ-2994 in a T-ALL animal model. Luciferase expressing RPMI8402 cells were injected via tail vein into irradiated NSG mice. Leukemia establishment was determined using bioluminescent imaging, and mice were randomized into two groups, vehicle versus RZ-2994 treatment once disease was established. Mice were treated with 100 mg/kg IP daily for 2 weeks and disease burden evaluated. The drug was well tolerated, with stable weights for both the vehicle and treatment cohorts (Fig. 5F). RZ-2994 treatment led to a decrease in leukemia burden in the bone marrow and spleen (Fig. 5G), supporting further compound optimization and pre-clinical evaluation of this pathway in T-ALL.

### SHMT inhibition is efficacious in the setting of methotrexate resistance

Methotrexate is a backbone of ALL chemotherapy treatment. Although testing of methotrexate sensitivity is not done routinely, ALL at the time of relapse has been shown to be relatively methotrexate resistant^48,49^. Although highly effective in upfront therapy, inhibitors of the one-carbon folate pathway are not typically used at the time of relapse. We thus addressed whether targeting of SHMT1 and SHMT2 with RZ-2994 can be effective in the setting of methotrexate resistance. We developed methotrexate resistant cell lines by growing PF382 and KOPTK1 cell lines in increasing concentrations of methotrexate over several months and found methotrexate resistant cell lines remained sensitive to RZ-2994 (Fig. 6A and 6B).

**Figure 6:**
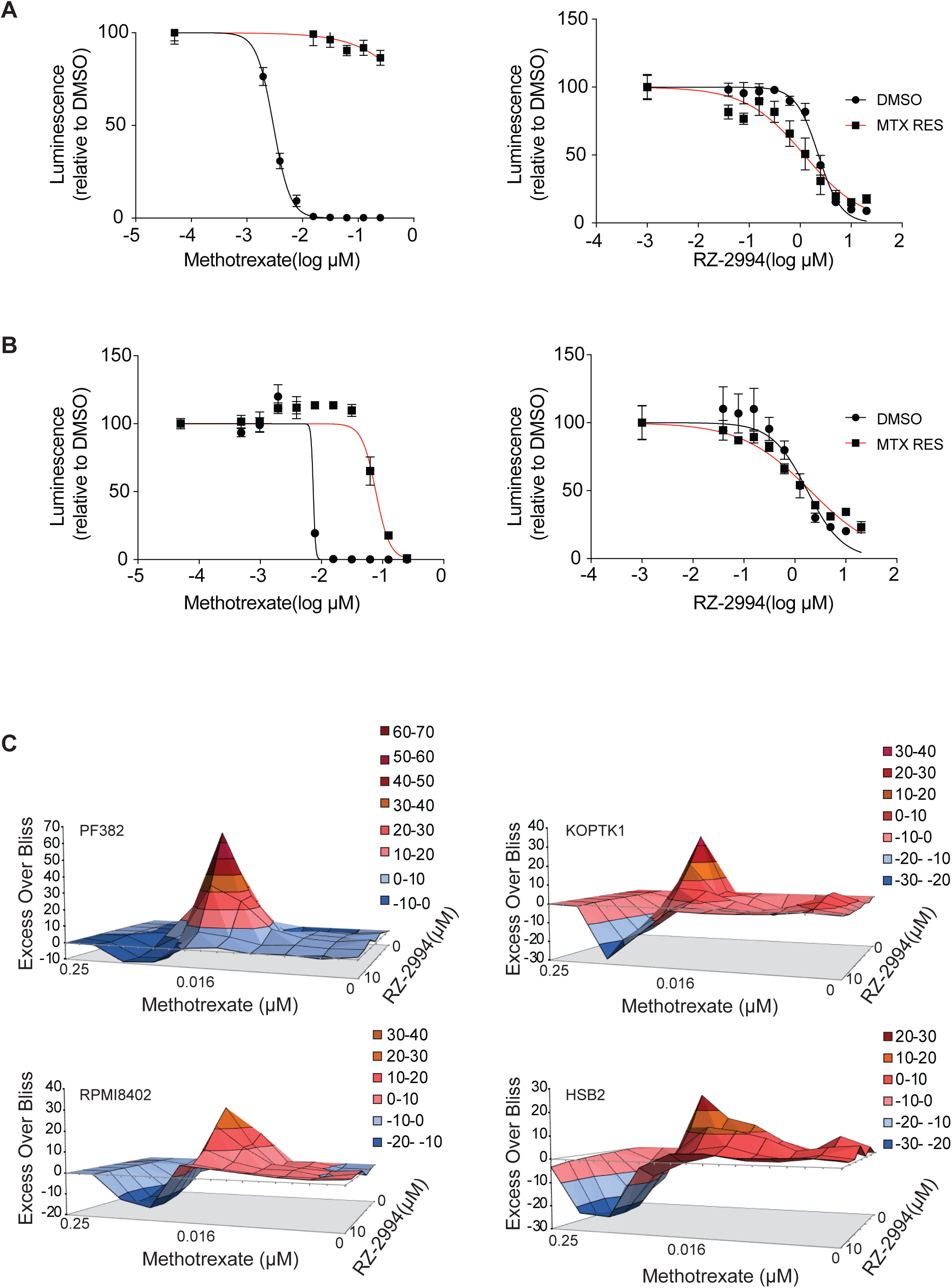
RZ-2994 is effective in the setting of methotrexate resistance. PF382 (A) and KOPTK1 (B) cells were grown to methotrexate resistance and then tested for sensitivity to RZ-2994. Parental and methotrexate resistant cells were tested with a range of RZ-2994 concentrations and viability evaluated at day 6 by an ATP-based assay as the percentage of viable cells relative to a DMSO control. Shown are the mean ± standard deviation (SD) of 4 replicates. (C) Excess over Bliss analysis for the combination of RZ-2994 with methotrexate in PF382, KOPTK1, RPMI8402 and HSB2 cells treated for 6 days in replicates of 4.

Targeting a pathway at two different nodes can be clinically efficacious as demonstrated using the combination of methotrexate with mercaptopurine for treatment of ALL. We thus tested the combination of methotrexate with RZ-2994. We treated the PF382, KOPTK1, RPMI8402 and HSB2 cells with RZ-2994 in combination with methotrexate concurrently across a range of drug concentrations in a serially 2-fold dilution. Cells were treated in 384-well format in quadruplicate for each drug concentration combination and viability was assessed after 3 and 6 days of treatment using the CellTiter-Glo ATP-based assay. We used the Bliss independence model to assess for synergy. Based on this model, we observed a mixed response. The combination of RZ-2994 with methotrexate was antagonistic at the highest concentration of methotrexate, including around the IC_50_, but showed synergy at lower concentrations of methotrexate (Fig.6C). Given this mixed concentration-dependent response, caution would be needed in bringing this combination into a clinical setting.

## Discussion

Despite significant progress in the treatment of pediatric ALL since the 1950s, leukemia still accounts for the second leading cause of cancer-related death in children. For pediatric and adult patients with relapsed T-ALL, treatment options are limited, with disease often resistant to chemotherapy at the time of relapse. Thus, alternative therapies are needed.

In this study, we identified the one-carbon folate pathway as an enriched pathway dependency in T-ALL. One-carbon folate metabolism is critical for nucleotide synthesis, support of cellular methylation reactions via methionine and s-adenosyl methionine (SAM) production, redox regulation and support of lipid metabolism^3^. The role of one-carbon folate metabolism in the mitochondrial compartment, and its potential contribution to cancer metabolic reprogramming, has only recently come to light with the discovery of a role for glycine, serine, and glutamine in oncogenesis^46,50-52^. Although cytoplasmic components of the one-carbon folate pathway have long been targeted for cancer therapy, classically with drugs such as methotrexate and mercaptopurine, targeting the mitochondrial proteins of this pathway has not been explored. Two such mitochondrial proteins, NAD-dependent mitochondrial methylenetetrahydrofolate dehydrogenase/cyclohydrolase (MTHFD2) and serine hydroxymethyltransferase 2 (SHMT2), are among the most differentially expressed metabolic enzymes in cancer cells compared to normal cells^53,54^. Both MTHFD2 and SHMT2 overexpression have been associated with tumor pathogenesis and poor survival^46,55,56^.

Although acute leukemia is highly proliferative, the effect of one-carbon folate pathway inhibition using a novel inhibitor of SHMT1 and SHMT2, RZ-2994, was greater in T-ALL compared to AML and B-ALL. Over 70% of T-ALL have mutations leading to activation of NOTCH1 signaling, and this is associated with MYC overexpression. MYC controls a number of critical metabolic processes, including the one-carbon folate pathway. In fact, inhibition of SHMT1 and SHMT2 recapitulated gene expression changes associated with MYC inhibition and may contribute to the differential sensitivity of T-ALL to one-carbon folate pathway inhibition.

The mechanistic role of SHMT1 and SHMT2 that is specific to T-ALL pathogenesis remains elusive. MTHFD2 is classically described as contributing to one-carbon unit production through conversion of serine to glycine by SHMT2 and production of formate as a product of MTHFD2 and MTHFD1L activity. The cytoplasmic arm of this pathway relies on SHMT1 and can also produce glycine and formate^47^ intermediates contributing to the synthesis of purines/pyrimidines, as well as to the methionine and glutathione cycles. The direction of one-carbon and electron flow through this pathway, and the contribution of the cytoplasmic versus mitochondrial pathways, have been debated and may be different in normal compared to cancer cells and under variable nutrient conditions^47,54^. In T-ALL cell lines, formate supplementation rescued the effects of dual SHMT1/SHMT2 inhibition, as well as the anti-proliferative effect of combined SHMT1/SHMT2 knockdown. This contrasts with data in DLBCL, where a defect in exogenous glycine import affects the formate’s ability to rescue the effects of SHMT inhibition^19^. Interestingly, there was no depletion of SAM and methionine with RZ-2994 treatment (Supplementary Fig. 5). Maddocks et al. showed that the one-carbon folate pathway does not contribute one-carbon units to methionine in the presence of methionine replete conditions in cells in culture^57^, likely explaining the lack of methionine depletion in RZ-2994 treated cells.

Chemotherapy resistance is a key factor in cancer treatment failure. Inhibitors of the one-carbon folate pathway are used for treatment of cancers including ALL, osteosarcoma, breast, lung and many others, though predictors of response are not evaluated pre-treatment. We showed that T-ALL cell lines that are resistant to methotrexate remain sensitive to another inhibitor of the one-carbon folate pathway. It is also possible that resistance to inhibition of the one-carbon folate pathway may be overcome with novel inhibitors of this pathway though further *in vivo* testing is necessary.

RZ-2994, a recently developed inhibitor of SHMT1 and SHMT2, was reported to have poor stability in liver microsome assays^19^. We performed a PK study, however, that showed a t_½_=5.9 hours. Further optimization to increase *in vivo* stability and prolong target engagement is likely necessary for improved *in vivo* efficacy. In addition, RZ-2994 causes cell cycle arrest. Active drug combinations with an SHMT1/2 inhibitor with other drugs that cause cell death may be necessary to maximize efficacy of this approach. Often new drugs are combined with standard chemotherapy for treatment of patients with leukemia. However, the combination of RZ-2994 with standard chemotherapy, which is most toxic to rapidly proliferating cells, may be antagonistic and careful testing of combinations will be necessary for clinical implementation of an optimized inhibitor.

In summary, the combination of unbiased genome-wide screening identifying the one-carbon folate pathway as a dependency in T-ALL, as well as the preclinical efficacy of SHMT1 and SHMT2 inhibition using both chemical and genetic approaches, support further optimization of SHMT1/2 inhibitors. Given the efficacy of other one-carbon folate pathway targeting drugs in cancer, novel inhibitors of this pathway are likely to be effective in other disease types, both at the time of diagnosis as well as at the time of relapse and development of drug resistance.

## Abbreviations used

T-ALL: T-cell acute lymphoblastic leukemia;
B-ALL: B-cell acute lymphoblastic leukemia;
AML: acute myeloid leukemia;
DHFR: dihydrofolate reductase;
ALL: acute lymphoblastic leukemia;
MTHFD2: methylenetetrahydrofolate dehydrogenase-cyclohydrolase 2;
KEGG: Kyoto Encyclopedia of Genes and Genomes;
GSEA: Gene Set Enrichment Analysis;
SHMT1: serine hydroxymethyltransferase 1;
SHMT2: serine hydroxymethyltransferase 2;
MTHFD1L: methylene tetrahydrofolate dehydrogenase 1-like;
LC-MS: liquid chromatography-mass spectrometry.

## Acknowledgements

We would like to thank Adam Friedman, Nello Mainolfi, Vipin Suri and Mark Manfredi of Raze Therapeutics for providing RZ-2994. This research was supported with grants from a Rally Foundation/Bear Necessities Collaborative Grant (KS), National Cancer Institute R35 CA210030 (KS), K08 CA222684 (YP), F31 CA236036 (FFD), R35 CA242379 (MGVH), Hyundai Hope on Wheels grant (YP), AIRC (n. 17107, GR), Cubans Curing Children’s Cancers (4C’s Fund) (KS), Children’s Leukemia Research Association (KS) and When Everyone Survives (KS). AP receives support from the ERC Starting program (758848) and is supported by the St. Louis Association for leukemia research. M.G.V.H. also acknowledges support from the MIT Center for Precision Cancer Medicine, the Ludwig Center at MIT, SU2C, and a faculty scholars award from HHMI.

## Supplementary Figure Legends

**Supplementary Figure 1:** Graphs showing distribution of the ssGSEA Z-scores for the purine and pyrimidine metabolism pathways across cancer cell lineages represented in the Avana 19Q4 data set. A) The purine metabolism pathway is significantly enriched in T-ALL vs non-T-ALL hematopoietic (**P < 0.01, Mann-Whitney test) and T-ALL vs solid tumor (***P < 0.001, Mann-Whitney test). B) The pyrimidine metabolism pathway is significantly enriched in T-ALL vs non-T-ALL hematopoietic (***P < 0.001, Mann-Whitney test) and T-ALL vs solid tumor (***P < 0.001, Mann-Whitney test).

**Supplementary Figure 2:** A) Heatmap of ssGSEA projection for the primary ALL dataset from Den Boer et al.^58^ on the collection of KEGG canonical pathways. T-ALL samples are highlighted in red. Graphs showing the distribution of the ssGSEA Z-scores for the one-carbon folate pathway in the GSE33315 (B) and GSE13351 (C) data sets (***P<0.001, Mann-Whitney test).

**Supplementary Figure 3:** Cell cycle analysis in T-ALL cells treated with increasing concentrations of RZ-2994.

**Supplementary Figure 4:** RNAseq was performed for KOPTK1 cell line treated with RZ-2994. Volcano plots showing quantitative comparison of gene sets from MSigDB v7.0 using ssGSEA. Volcano plots compare DMSO versus RZ-2994 after 1 day of treatment. All datasets above dashed red line have P-value ≤ 0.05. Gene expression changes associated with 1-day RZ-2994 treatment show enrichment for A) amino acid metabolism, B) MYC targets and C) cell cycle pathways. Top scoring GSEA plots are shown below the associated volcano plots.

**Supplementary Figure 5:** Heatmaps showing metabolites that were significantly changed with RZ-2994 treatment across 3 cell lines. Cell lines were treated with 2 µM RZ-2994 for 3 days, metabolites extracted and profiled using LC-MS. Raw peak areas were normalized to internal standards. Heatmap shows normalization of the relative metabolite abundance per metabolite.

**Supplementary Figure 6:** Volcano plots showing effect size for *SHMT1* or *SHMT2* knockout in 689 cell lines in the Avana 19Q4 dataset. The differential dependency gene level scores for the T-ALL lineage were determined for T-ALL vs. all other non T-ALL cell lines (A), and also for T-ALL vs. all other non T-ALL hematopoietic cell lines (B), in order to eliminate the bias induced by the hematopoietic lineage. C) Graph showing CERES dependency score for *SHMT1* knockout in 689 cancer cell lines screened as part of the Avana 19Q4 data set. T-ALL cell lines are indicated in red. Dotted lines signify level of significant dependency (CERES score<-0.5) or proliferative advantage (CERES score >0.5). D) Graph showing CERES dependency score for *SHMT2* knockout in 689 cancer cell lines screened as part of the Avana 19Q4 data set. T-ALL cell lines are indicated in red.

