## Supplementary figures and images for "Targeting serine hydroxymethyltransferases 1 and 2 for T-cell acute lymphoblastic leukemia therapy"

### Supplemental figure 1

**A**

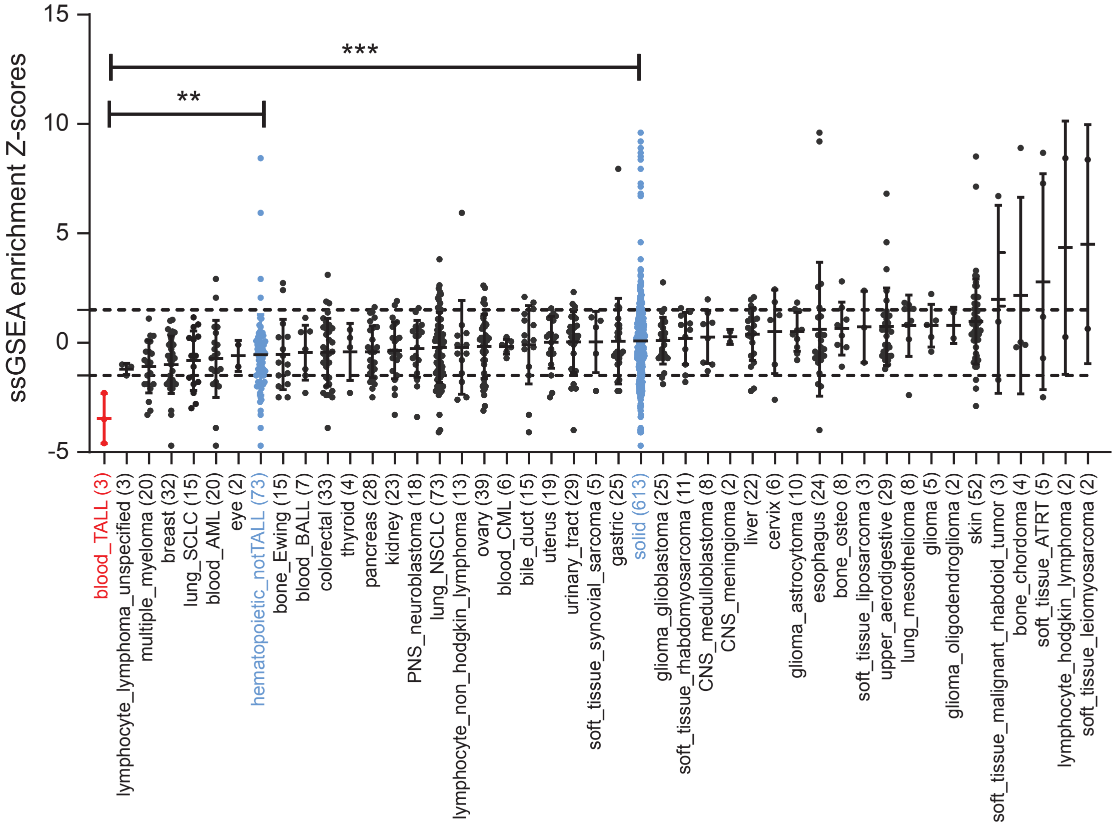


**B**

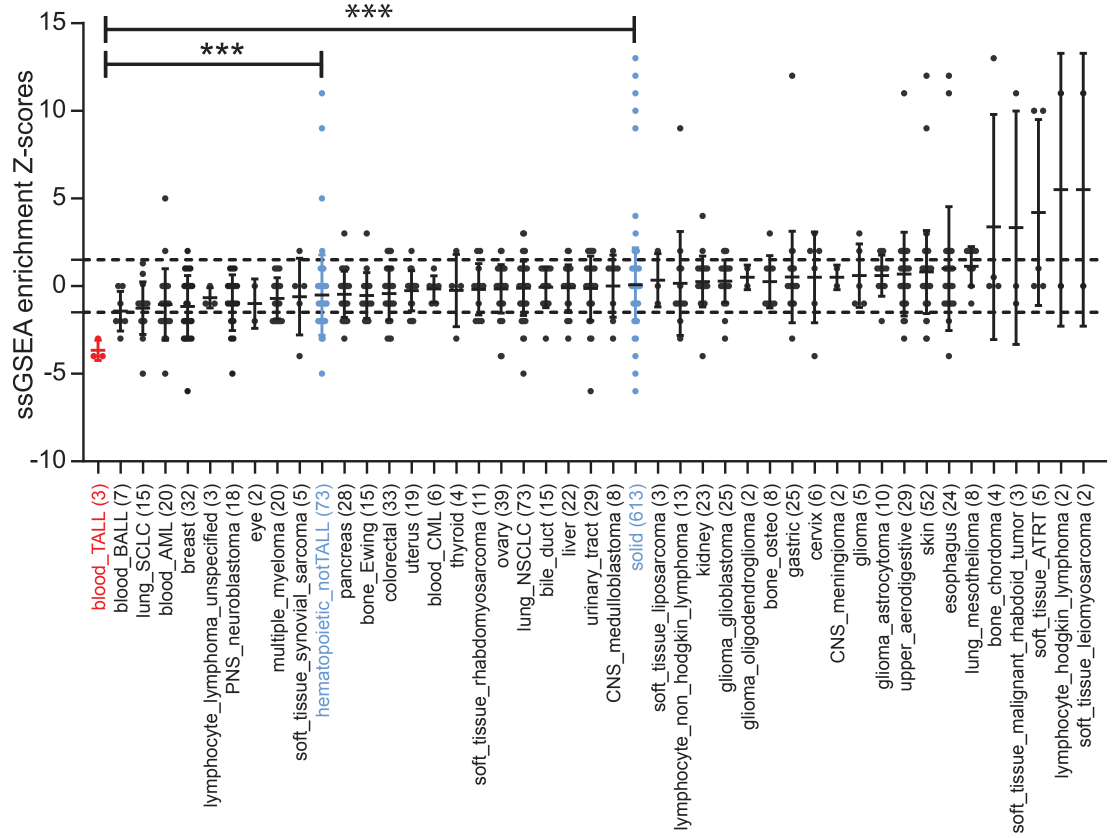


**Supplementary Fig. 1**

### Supplemental figure 2

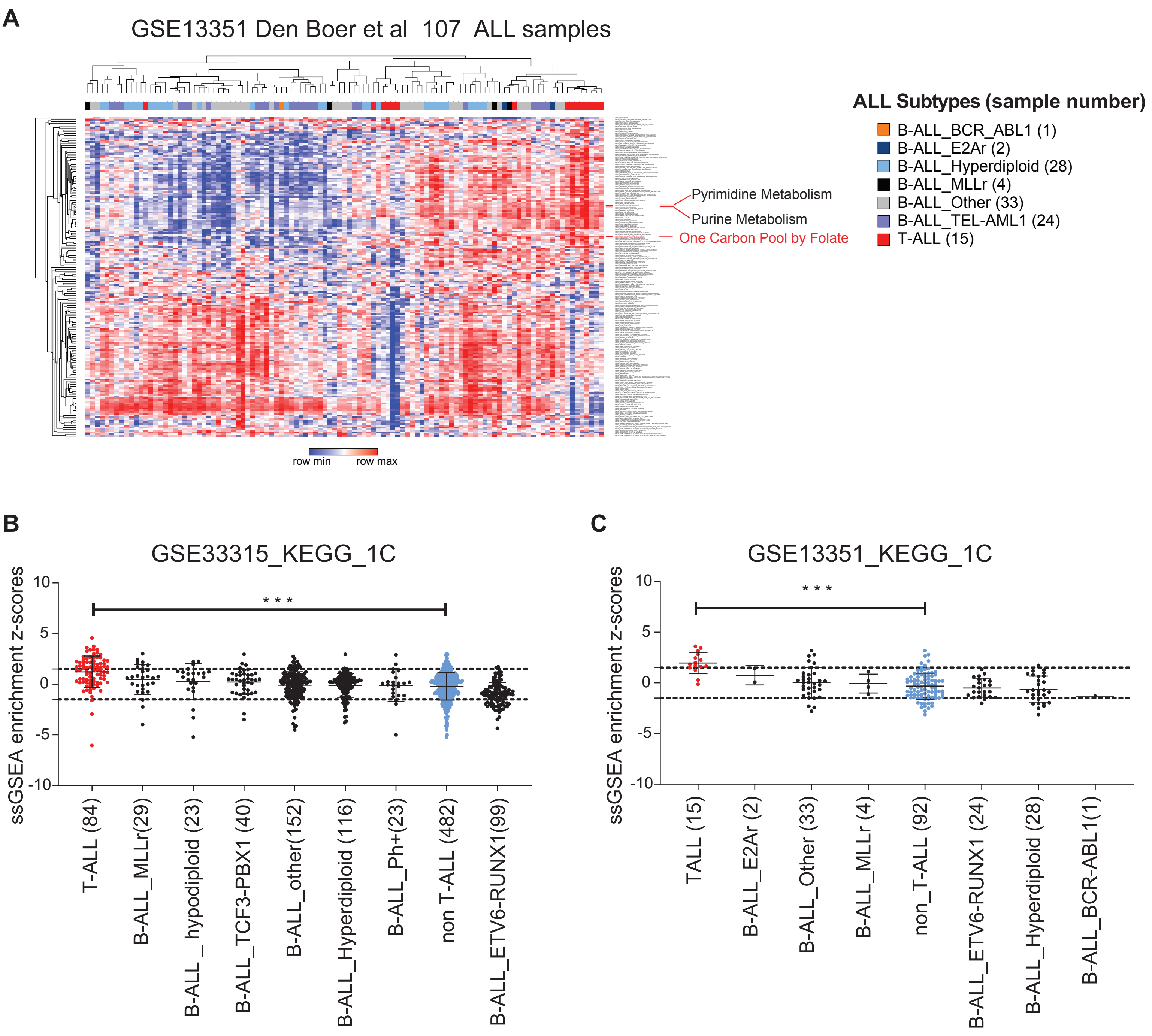


**Supplementary Fig. 2**

### Supplemental figure 3

A

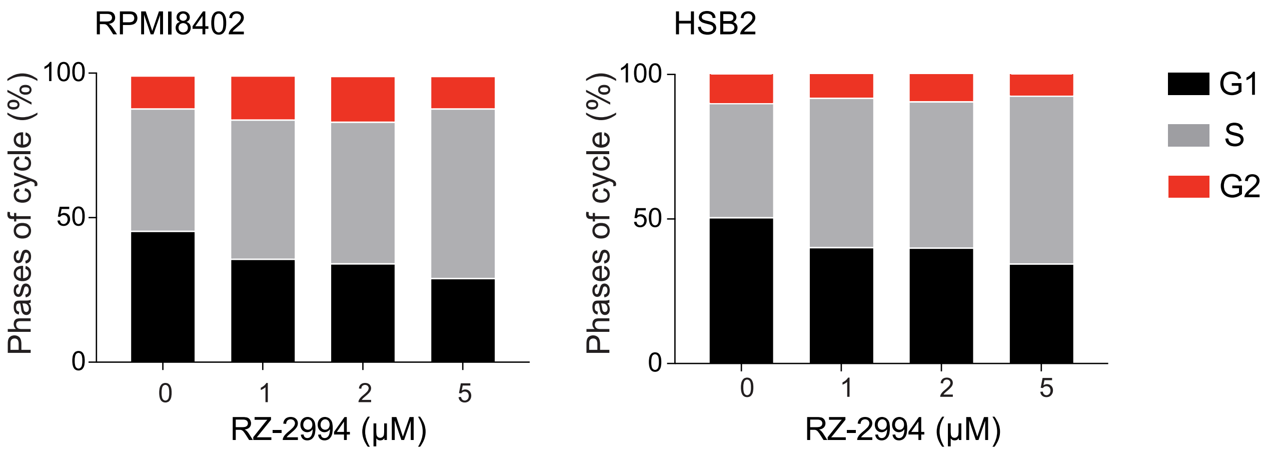


**Supplementary Fig. 3**

### Supplemental figure 4

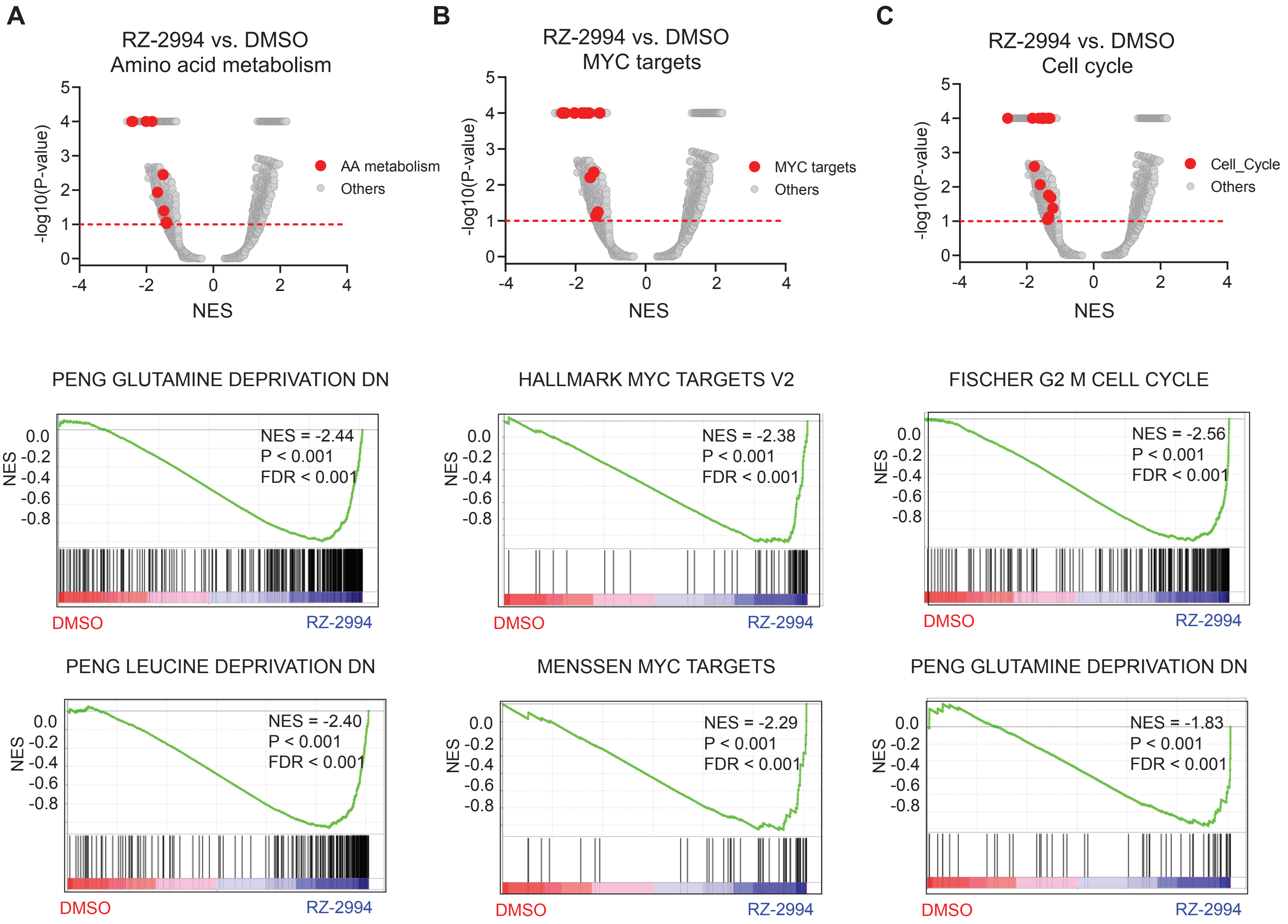


**Supplementary Fig. 4**

### Supplemental figure 5

**A**

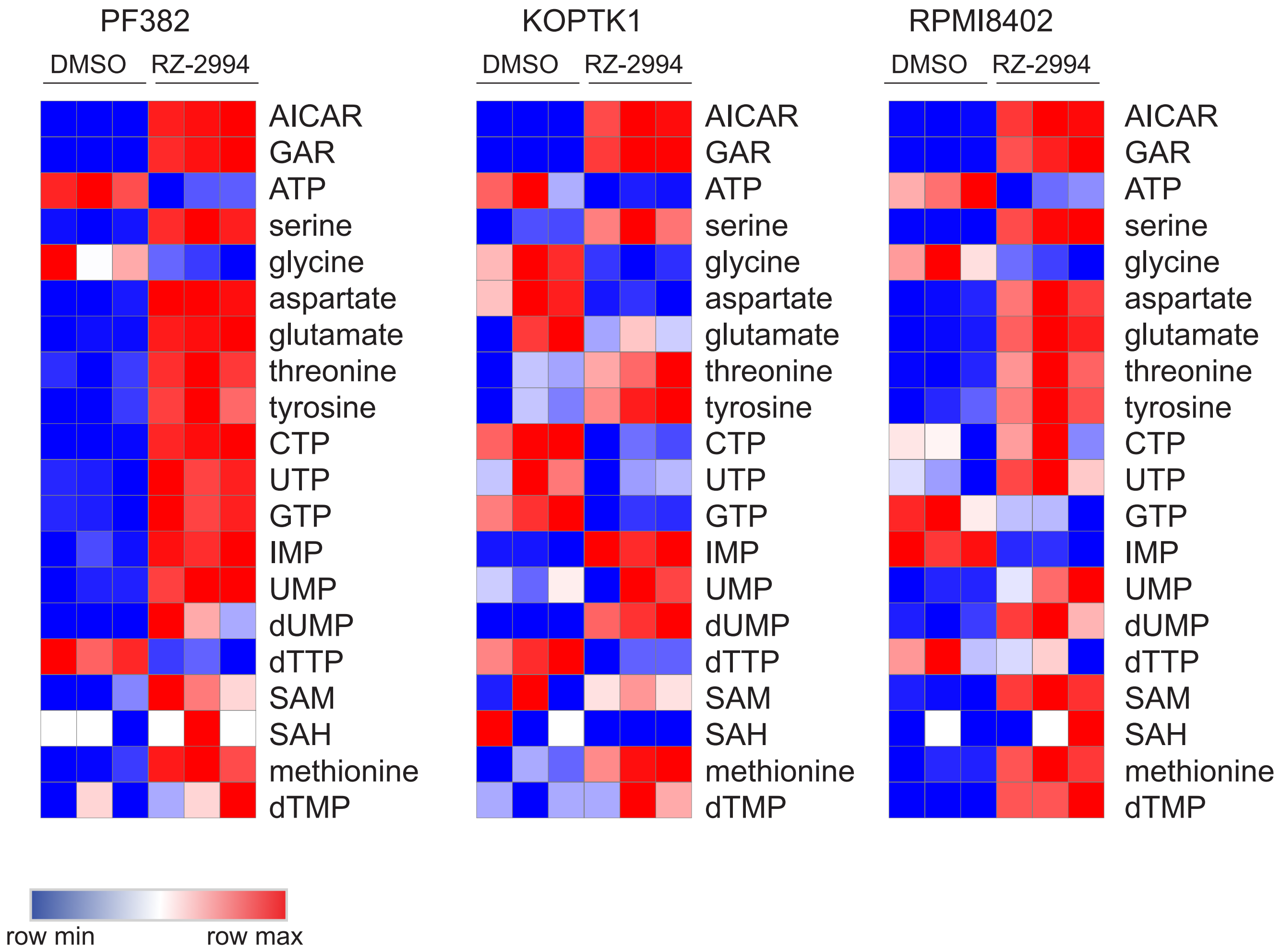


**Supplementary Fig. 5**

### Supplemental figure 6

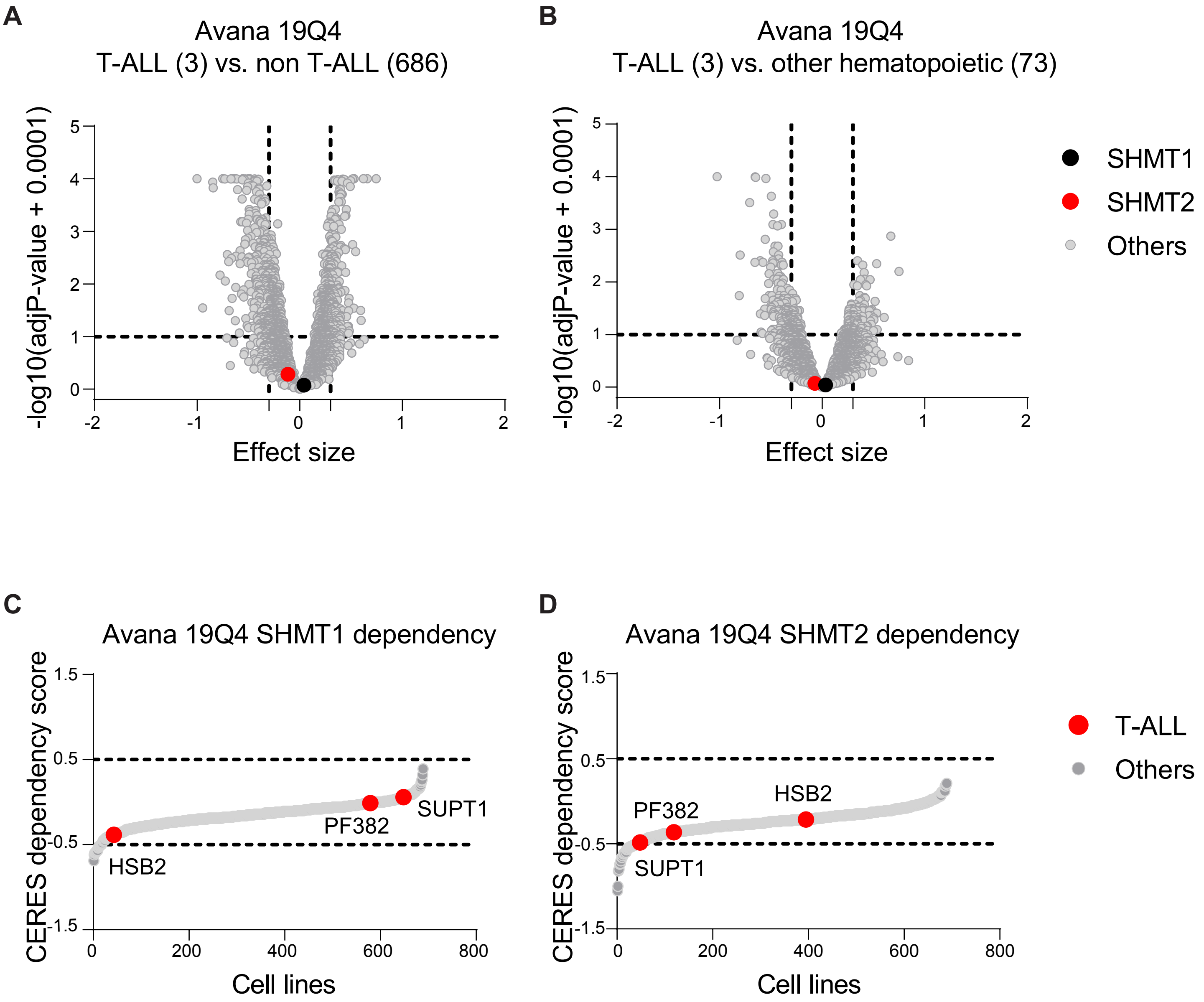


**Supplementary Fig. 6**
